# Characterization of miRNAs encoded by Autographa californica nucleopolyhedrovirus

**DOI:** 10.1101/2020.05.13.094193

**Authors:** Jinwen Wang, Ke Xing, Peiwen Xiong, Hai Liang, Mengxiao Zhu, Shaojuan Huang, Jin Zhao, Xinghua Yu, Xiaolian Ning, Runcai Li, Xunzhang Wang

**Affiliations:** School of Life Sciences, Sun Yat-sen University, Guangzhou 510275, China; School of Life Sciences, Guangzhou University, Guangzhou 510006, China

**Author notes:** Corresponding authors: Jinwen Wang and Xunzhang Wang.

**Keywords:** Baculovirus AcMNPV, miRNA, Target, Regulation, Host

## Abstract

Two Autographa californica nucleopolyhedrovirus (AcMNPV) encoded miRNAs, AcMNPV-miR-1 and -miR-3, have been reported in 2013 and 2019. Here, we present an integrated investigation of AcMNPV-encoded miRNAs. Six candidate miRNAs were predicted through small RNA deep sequencing and bioinformatics, of which, five validated by experiments. Three miRNAs perfectly matched the coding sequence of viral genes. The other two are located in coding sequences of viral genes. Targets in both virus and host were predicted and subsequently tested using dual-luciferase reporter assay. The validated targets were found mainly in AcMNPV, except for the targets of AcMNPV-miR-4, which are all host genes. Based on reporter assays, the five miRNAs predominantly function by down-regulating their targets, though individual target is slightly up-regulated. The transcription start sites of these miRNAs were analyzed. Our results suggest that AcMNPV-encoded miRNAs function as fine modulators of the interactions between host and virus by regulating viral and/or host genes.

**Author summary:** Virus-encoded miRNAs have been widely reported as modulators participating in almost all biological processes. However, among *Baculoviridae*, which consists of a large family of dsDNA viruses that infect numerous beneficial insects and agricultural pests, only several have been reported encoding miRNAs. To clarify the roles of AcMNPV-encoded miRNAs in host–pathogen interactions, we employed RNA deep sequencing and series of experimental approaches identifying AcMNPV-encoded miRNAs, followed by target validation and function deduction. Among them, AcMNPV-miR-1 and AcMNPV–miR-3 have been reported in 2013 and 2019, respectively. This study reveals the sites of these miRNAs in the genome, both in coding sequences and complements, suggesting diverse functions. These miRNAs target genes in the virus itself and in the host, largely by suppressing expression, with some enhancing it. The transcription initiations of the miRNAs were analyzed. Our results provide some insight into the finely regulated process of baculovirus infection.

## Introduction

MicroRNAs (miRNAs) are small, noncoding RNAs ∼22 nucleotides in length derived from primary transcripts (pri-miRNAs). They undergo endonucleolytic processing twice, becoming precursor miRNAs (pre-miRNAs) and then final, mature miRNAs that target messenger RNAs (mRNAs) via base pairing. miRNAs modulate the stability and/or translational potential of their mRNA targets to achieve regulatory control of virtually every biological process [1-3]. In many cases, a single miRNA can regulate many targets, and one mRNA can interact with multiple miRNAs [2, 3]. Therefore, the network between miRNAs and their target mRNAs is comprehensive and complex.

In the last decade, many virus-encoded miRNAs have been identified, and their targets and functions have been characterized and deduced. According to these reports, viral miRNAs are involved in multiple cellular processes, such as immunity, development, differentiation, apoptosis, and proliferation [2, 3]. In early 2005, SV miRNAs encoded by SV40 were reported to reduce the expression of viral T antigens to facilitate successful virus infection [4]. Transcriptional regulation of three primary miRNAs in herpes simplex virus 2 may play an important role in the switch between latent and productive infection [5]. The insect virus-encoded miRNA HvAV-miR-1, which is derived from the major capsid protein gene, down-regulates expression of viral DNA polymerase I and hence inhibits viral replication [6]. Although evidence to date indicates that virus-encoded miRNAs have important functions in virus-host interactions, a deep and detailed understanding of most virus-encoded miRNAs is still lacking.

*Baculoviridae* comprises a family of large, circular, double-stranded DNA viruses that are pathogens of arthropods. To date, three types of alpha baculoviruses, Bombyx mori nucleopolyhedrovirus (BmNPV), AcMNPV and Spodoptera litura nucleopolyhedrovirus (SpltNPV), have been reported to encode miRNAs [7-9]. Four BmNPV-encoded miRNAs were reported in 2010 [7], two of which were later functionally characterized. BmNPV-miR-1 inhibits expression of host exportin-5 cofactor, and Ran is mainly involved in small-RNA transport from the nucleus to the cytoplasm [10]. BmNPV-miR-3 regulates expression of DNA-binding proteins (P6.9) and other late genes, which are vital for the late stage of viral infection in the host *B. mori* [11]. Ten SpltNPV miRNAs that were reported in 2015 have also been validated [9]. AcMNPV, which is approximately 133 kb, serves as a model system for studying baculovirus molecular biology, was also reported encoding miRNAs, AcMNPV-miR-1 and AcMNPV-miR-3 in 2013 [8, 12] by us, respectively. AcMNPV-miR-1 can down-regulate viral gene *ac94*, reducing the production of infectious budded virions (BVs) and accelerating the formation of occlusion-derived virions (ODVs) [8]. In 2016, we further reported two more target genes (*ac95* and *ac18*) of AcMNPV-miR-1, and depicted the function pathway of this miRNA [13]. As for AcMNPV-miR-3, it mainly down-regulates viral gene *ac101*, meanwhile, it also regulates other five viral genes [12]. All these findings indicate that baculovirus-encoded miRNAs play roles in multiple aspects of virus infection.

Here, we provide an integrated characterization of these miRNAs which AcMNPV encoded. The investigation for AcMNPV-encoded miRNAs and their functions began with sequencing four small RNA samples collected from Sf9 cells infected with AcMNPV at three different time intervals and mock-infected in 2009. Subsequently, six viral candidate miRNAs were predicted by computational analysis, and five of which were validated by experiments. Targets were predicted by bioinformatics, some of which were validated experimentally.

## Materials and Methods

### Viruses and cell lines

*Spodoptera frugiperda* (*S. frugiperda*) IPLB-Sf21-AE clonal isolate 9 (Sf9) insect cells were cultured at 27°C in Grace’s medium (Gibco) containing 10% fetal bovine serum (Gibco). Hemolymph was extracted from infected fifth-instar *T. ni* larvae and used to infect Sf9 cells after passing through a 0.22-μm Millipore filter. The third-generation supernatant from infected Sf9 cells was used as a working AcMNPV stock. The viral inoculum was allowed to adsorb to cells for 1 hour of infection or 4 hours of transfection at 27°C. Time zero was defined as the time when the viral inoculum was replaced.

### RNA sample preparation and sequencing

Four independent total RNA samples were extracted from Sf9 cells infected with AcMNPV (10 PFU/cell) at 6, 12, and 24 h post-infection (p.i.) and mock-infected using TRIzol reagent (Invitrogen) according to the manufacturer’s protocol. RNA concentrations were measured using a spectrophotometer, and integrity was ensured through analysis of RNA on a 1% (w/v) agarose gel and using a bioanalyser. The RNA samples were sent to Beijing Genomics Institute ShenZhen Co. (BGI-ShenZhen) for small RNA sequencing. Small RNA libraries were generated based on a standard procedure [1]. Briefly, small RNA molecules were isolated by size fractionation on a 15% polyacrylamide gel, RNA adapter was ligated to the 3’ end, reverse transcriptase was used to generate first-strand cDNA, followed by PCR amplification of the cDNA. Size fractionation of the cDNA was performed, followed by Solexa sequencing. Ultimately, these RNA-Sequence data were submitted to NCBI SRA database, the accession number is PRJNA523298.

### Prediction of viral miRNA

To filter data from raw sequences, reads without 3’ adaptors and reads with less than 18 nt were discarded. Clean sequences of 18 to 30 nt were considered to be credible reads. Four small RNA libraries were generated, each containing more than 1 million unique reads. These reads were identified by scanning the AcMNPV genome (accession number NC_001623.1) in both orientations for sequences of 50 to 120 nt in length and could be folded into stem-loop structures. These candidate segments were validated using two online Web servers: MiPred and CSHMM. The sequenced reads were mapped to the candidate segments using the SOAP program. Candidates without matched reads were removed.

### Northern blot analysis

Total RNA was extracted from Sf9 cells infected with AcMNPV at different time intervals and mock-infected using TRIzol reagent (Invitrogen) according to the manufacturer’s protocol. 15 micrograms of each total RNA samples, as well as siRNA ladder marker (Takara), were separated on a 15% denaturing polyacrylamide gel electrophoresis (PAGE) (acrylamide : bis-acrylamide ratio, 19 : 1). RNAs were then transferred to a positively charged nylon membrane (Roche) using a semi-dry transfer cell (Trans-blot®SD Bio-Rad) and UV cross-linked (HL-2000 HybriLinker, UVP) to the membrane. The reverse complementary RNA probes were 5’ end-labeled with digoxin (TaKaRa). Hybridizations and washes were carried out using DIG Block and Wash Buffer (Roche). Signals were detected using CSPD (Roche) through GBOX-CHEMI-XT4 (Syngene, Gene Company Limited) or Tanon-5200 Chemiluminescent Imaging System (Tanon Science & Technology).

### Poly (A)-tailed RT-PCR

Small RNAs were prepared using a miRNA isolation kit (OMEGA), and poly (A) polymerase was used to add poly (A) tails to the 3’ ends of the small RNAs. miRNA PrimeScript RT Enzyme and a unique oligo-dT adaptor primer (including a universal primer) were used to transcribe poly (A)-tailed miRNAs to generate a cDNA library according to the PrimeScript™ miRNA RT-PCR Kit (Takara) instructions. PCR was performed on the cDNA using a miRNA-specific forward primer and adaptor-specific reverse primer (universal primer).

### Stem-loop RT-PCR

Total RNA was prepared as described above. cDNA was synthesized from the total RNA using 6-8 nucleotide-long miRNA-specific stem-loop primers, as previously described, per the manufacturer’s instructions for PrimeScript™ miRNA RT-PCR Kit (Takara). The PCR mixture contained 3 μl cDNA, 3 μl 10× PCR buffer, 1 μl 10 mM dNTPs, 1 μl each 10 μM miRNA specific forward and universal reverse primers, 0.5 μl 5 U/μl Taq polymerase (Takara) and nuclease-free water to 30 μl. The PCR reaction was incubated at 94°C for 3 min, followed by 30 cycles of 94°C for 30 s, 55°C for 30 s, 72°C for 30 s, and a final extension at 72°C for 8 min.

### qRT-PCR detection of miRNA expression

RNA samples were extracted using a miRNA Isolation Kit (Ambion) and reverse transcribed using a PrimeScript™ RT reagent kit (TaKaRa). The reverse transcription product was used to perform quantitative PCR (qPCR) with a real-time PCR detection system (LightCycler®480; Roche). The 10-μl qPCR reaction mixture contained 1 μl cDNA, 5 μl 2×SYBR®Premix Ex Taq™ II (TaKaRa), 0.75 μM primers and nuclease-free water to 20 μl. PCR was conducted in the LightCycler®480 system (Roche) with a denaturation program of 8 min at 95°C followed by 40 cycles at 95°C for 10 s and 60°C for 30 s. Then, a program of 95°C for 1 s, 65°C for 15 s and 95°C continuously was used to obtain melting curves. The quantitative reverse transcription-polymerase chain reaction (qRT-PCR) data for each sample was calculated using the 2^-ΔΔ*CT*^ (where *C*_*T*_ is the threshold cycle) method [14]. A 5S rRNA reverse transcript generated using random primers (TaKaRa) was used as a reference. Each reaction was performed in triplicate in at least three independent experiments.

### Target prediction

All transcript sequences of AcMNPV, genome and transcriptome data of *S. frugiperda* and *Bombyx mori* (*B. mori*) were downloaded from the NCBI database and searched for potential targets of the predicted virus miRNAs using the miRanda program. The parameters assigned for miRanda were a Smith-Waterman hybridization alignment score greater than 90 (Smith and Waterman, 1981), a ΔG of miRNA::mRNA duplex less than -20 kcal/mol, and a scaling factor equal 2, with all other parameters set as default.

### Dual-luciferase reporter assay

The predicted binding site of each candidate target gene was amplified using corresponding primers. The PCR product was inserted into the multiple cloning site (MCS) of the psicheck-2 vector (supplied by Promega) downstream of the *Renilla* luciferase gene according to the manufacturer’s instructions. Selected plasmids with binding-site fragments were sequenced and then quantified using a NanoDrop 2000c spectrophotometer (Thermo Scientific). The mutational sequence of each binding site was synthesized by BGI-ShenZhen and cloned into the psicheck-2 vector using the method described above. Mutations were confirmed by sequencing. Using the above strategies, two types of plasmids were constructed for each possible target gene. An unrelated target sequence (*B. mori* actin 3) was inserted into the same vector as a universal control for all genes. Cel-miR-239b-5p, a miRNA from *Caenorhabditis elegans*, was used as the miRNA control. More details can refer to our previous papers [13].

## Results

### Computer prediction of miRNAs from AcMNPV

To search for AcMNPV-encoded miRNAs, four independent small RNA read libraries from AcMNPV-infected and mock-infected Sf9 cells at different time intervals were generated through deep sequencing. Each RNA library contained more than 1 million unique reads, which were further sorted by employing many credible filters based on published work [1, 15]. These reads were then mapped onto the AcMNPV genome (accession number NC_001623.1) in both orientations to search for sequences of 50 to 120 nt in length with characteristics that could be folded into a stem-loop structure, the typical structure of miRNA precursors. The minimum free energy (MFE) of the stem-loop structures was further determined using the MFold program. Ultimately, six small RNAs met all the criteria, and those qualified by manual checking were considered candidate AcMNPV miRNAs (Fig 1) (Table1). Three candidate miRNAs are located in the complimentary strands of viral genes. MR147 perfectly matches a segment in the coding sequence of the viral gene *ac94* (ODV-E25); MR40 and MD150 complement the coding sequence of the viral genes *lef-6* and *ac101* (BV/ODV-C42), respectively (Fig 2A-C) (Table1). The other three candidates are located in the coding sequence (CDS) of viral genes. MR139 is in the 5’ region of *ac-cg30*, MD222 is in the 5’ region of *ac-49k* and *ie-01*, and MD190 is in the 3’ region of *ac-v-cath* (Fig 2D-F) (Table1). The two types of distribution of the candidate miRNAs might suggest different functions and effects.

**Table 1.**
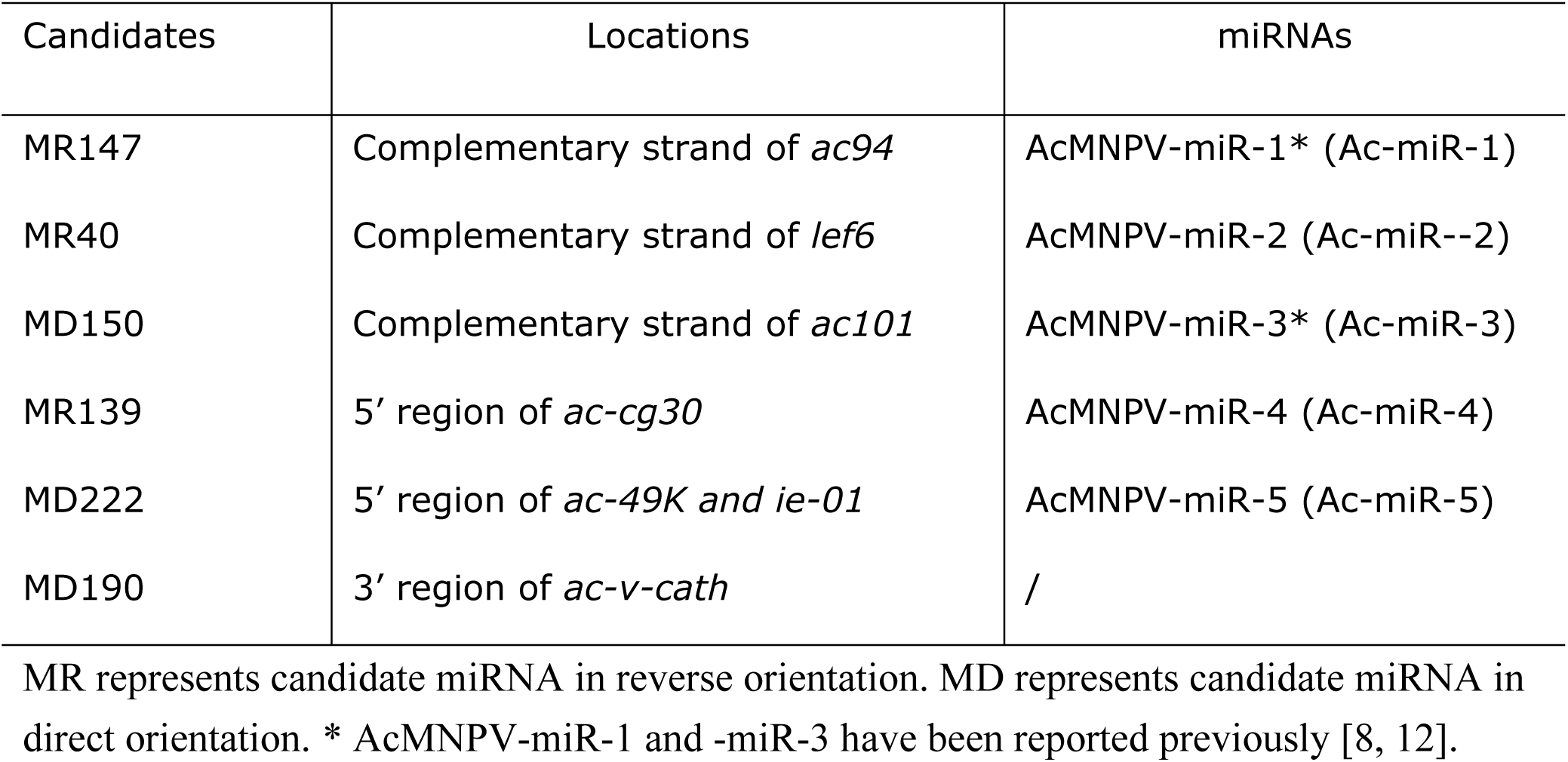
The predicted and validated miRNAs.

**Fig 1.**
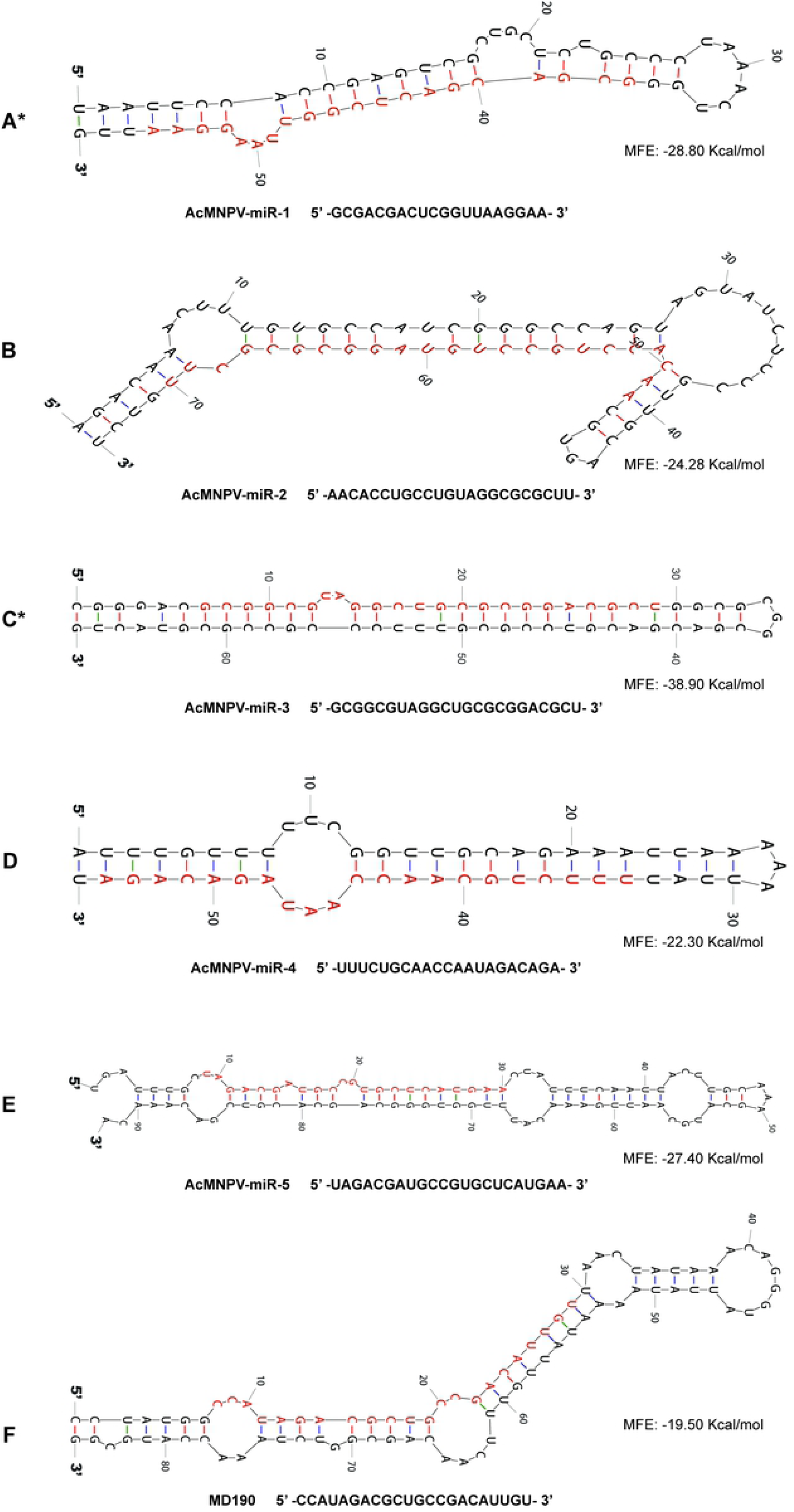
Secondary structures of the predicted AcMNPV encoded miRNA precursors and sequences of mature miRNAs. Secondary structures generated via the Mfold program (http://unafold.rna.albany.edu/?q=mfold/RNA-Folding-Form). The matured miRNA sequences are shown in red. MFE: minimum free energy. * The results of AcMNPV-miR-1 and -miR-3 are the same as our previous reports [8, 12].

**Fig 2.**
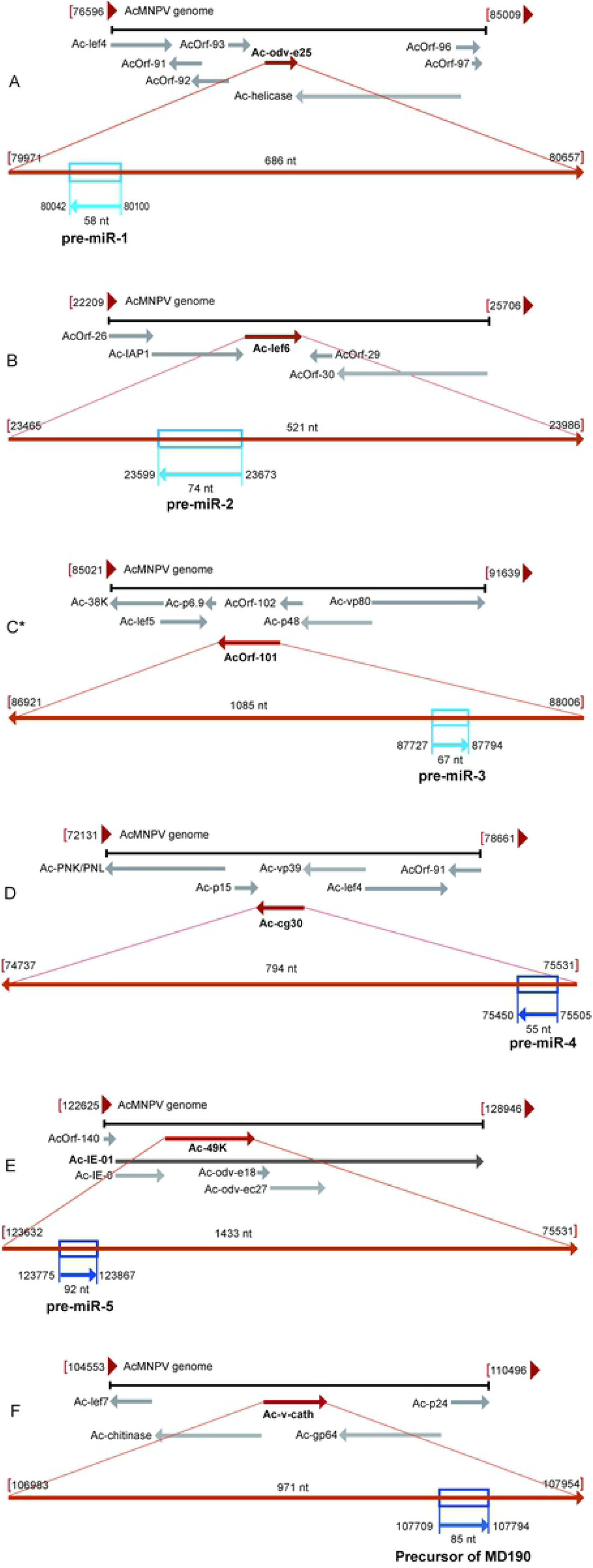
Genomic location and orientation of the predicted AcMNPV encoded miRNA precursors. The precursors, located genes and adjacent genes, and their orientation on the viral genome are represented in blue, red and light grey arrow lines, respectively. Light blue for miRNAs located in complementary strand and dark blue for those in coding strand. * AcMNPV-miR-3 has been reported previously [12].

### Expression validation of predicted candidate miRNAs

We employed several different experiments to assess expression of the predicted candidates. RNA samples were isolated from Sf9 cells with or without AcMNPV infection, and cDNAs were prepared as described in Materials and Methods.

First, poly (A)-tailed RT-PCR and stem-loop RT-PCR were carried out to detect predicted miRNA expression, and the expression of all six candidates was double-confirmed by sequencing of the PCR products from the two approaches (data not shown).

Then, northern blot hybridization (Fig 3) was performed with complementary RNAs that were end-labeled with digoxin specific for each predicted miRNA, followed by qRT-PCR (Fig 4) using stem-loop primers. The expression of candidate MR147 was further validated by northern blotting and qRT-PCR, as the first one, designated as AcMNPV-miR-1, which was reported in 2013 [8]. Candidate MR40 can be detected obviously at 3 h p.i. by northern blotting and qRT-PCR, distinct blotting bands also exhibited at 6, 12, 24 and 48 h p.i., and qRT-PCR showed the maximum expression at 6 h p.i. (Fig 3 and 4A), hence, designated as AcMNPV-miR-2. The expression of MD150 was also validated by both northern blotting and qRT-PCR, and designated as AcMNPV-miR-3, reported in 2019 [12]. In that report [12], AcMNPV-miR-3 can be detected obviously at 6 h p.i. To make clear if this miRNA expresses more earlier, we re-conducted northern blotting and qRT-PCR, and found AcMNPV-miR-3 can be detected distinctively at 3 h p.i. (Fig 3 and 4B). With regard to candidate MR139, distinct bands appeared at 24 h p.i. and maintained a similar signal intensity until 48 h p.i., qRT-PCR can detect it at 12 h p.i. and its maximum expression at 24 h p.i. (Fig 3 and 4C), then, it is designated as AcMNPV-miR-4. For candidate MD222, a 22-nt blotting band appeared at 3 h p.i., and the signals increased with time, qRT-PCR detected the obvious expression at 12 h p.i., increasing to a maximum at 24 h p.i., afterwards, the expression levels decreased at 36 and 48 h p.i. (Fig 3 and 4D), it is designated as AcMNPV-miR-5 (Table1). As for candidate MD190, qRT-PCR detected the similar expression pattern as that of AcMNPV-miR-5, but repeated northern experiments showed a 22-nt aim-sized blotting band in mock-infected sample lane, besides in each virus-infected sample lanes (3, 6, 12, 24, and 48 h p.i.) (data not shown), means MD190 may not be a true miRNA. In addition, precursors of above miRNAs were also blotted (Fig 3).

**Fig 3.**
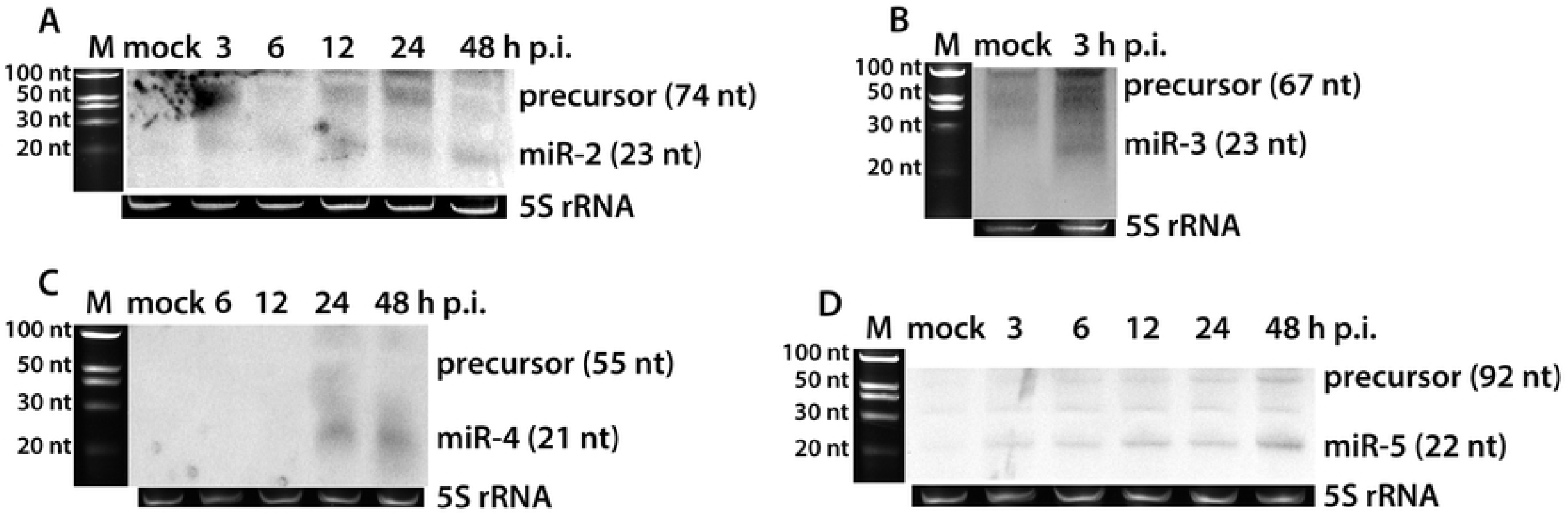
Northern blot confirmation of the expression of predicted miRNAs. A to D, MR40 (AcMNPV-miR-2), MD150 (AcMNPV-miR-3), MR139 (AcMNPV-miR-4), MD222 (AcMNPV-miR-5), respectively. Lane 1, RNA isolated from mock-infected Sf9 cells; lane 2 to 6, RNA isolated from Sf9 cells infected with AcMNPV at different time intervals. 5S rRNA, used as a loading control, and a siRNA ladder marker (TaKaRa) were detected under UV light before transfer.

**Fig 4.**
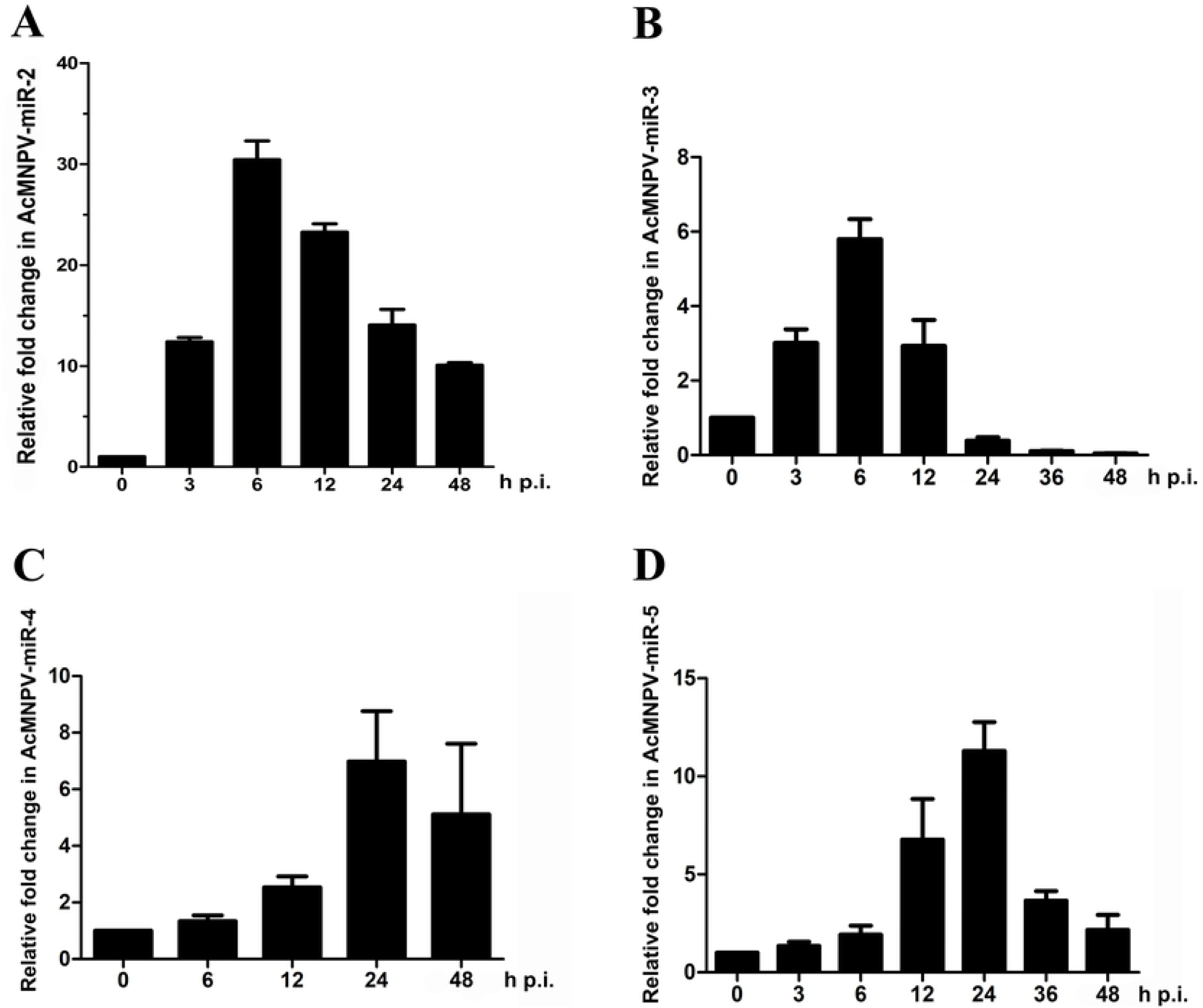
Relative expressions of predicted miRNAs determined by qRT-PCR. A to D, AcMNPV-miR-2 to AcMNPV-miR-5, respectively. Columns represent fold changes in miRNA expression normalized to 5S rRNA expression. RNA samples were isolated from Sf9 cells infected with AcMNPV at different time intervals, and the 2^-ΔΔCt^ method was used to calculate the RNA level in each sample. The Cq value of the no template control (NTC) was >40. Each reaction was performed in triplicate in at least three independent experiments. The error bars stand for the standard deviation.

Through the above four experimental approaches, Five of the six predicted candidate miRNAs were confirmed as true miRNAs encoded by AcMNPV.

### Target prediction and verification

miRNAs carry out their functions through their targets, and for viral miRNAs, their targets can be the virus’ own genes as well as the host’s genes. miRNAs can bind to the 3’ or 5’ UTR of the target mRNA, and they also can bind to coding sequences to manipulate target gene expression. To find potential targets of the five miRNAs, transcript sequences of AcMNPV and host (*S. frugiperda* and *B. mori*) were examined using the MiRanda program. Verification of the candidate target genes was achieved using a dual-luciferase reporter assay, which detects target expression using the *Renilla* luciferase gene as the reporter and the *firefly* luciferase gene as the reference. To verify a candidate target as a true target of one of the miRNAs, the recombinant plasmid psiCHECK-target was constructed by inserting the predicted miRNA binding site region into the psiCHECK-2 vector downstream of the *Renilla* luciferase gene. We also constructed a psiCHECK-target-mut plasmid with mutated binding sites and a psiCHECK-A3 plasmid with the *B. mori* actin 3 CDS inserted into the psiCHECK-2 vector as a control (Fig 5) [8, 13].

**Fig 5.**
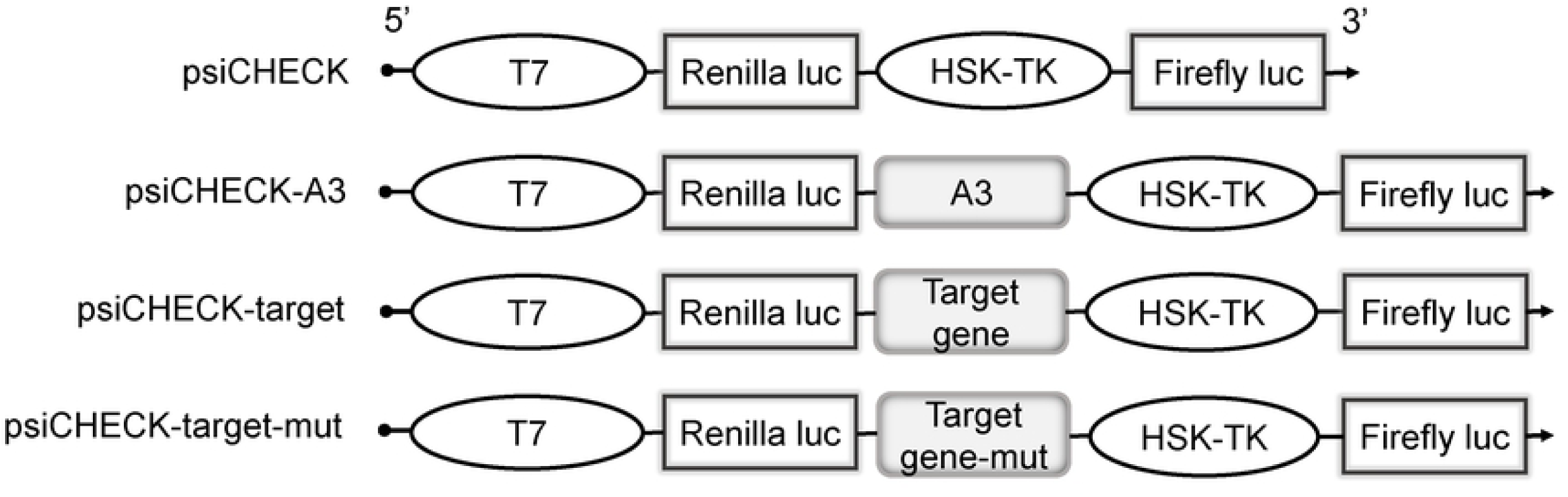
Schematic of luciferase reporter constructs for target gene verification. The predicted target genes CDS, mutated target binding site sequence, and CDS of an unrelated target *Bombyx mori* actin 3 inserted downstream of the *Renilla* luciferase (*luc*) gene, respectively.

Eight candidate target genes for AcMNPV-miR-1 were generated through above prediction analysis approaches. Among them, dual-luciferase reporter assay showed AcMNPV-miR-1 efficiently down-regulated *Renilla* luciferase activity due to perfect complementation with the *ac94* (*odv-e25*) CDS, moderately down-regulated *ac95* and up-regulated *ac18* (Table 2) [8, 13].

**Table 2.**
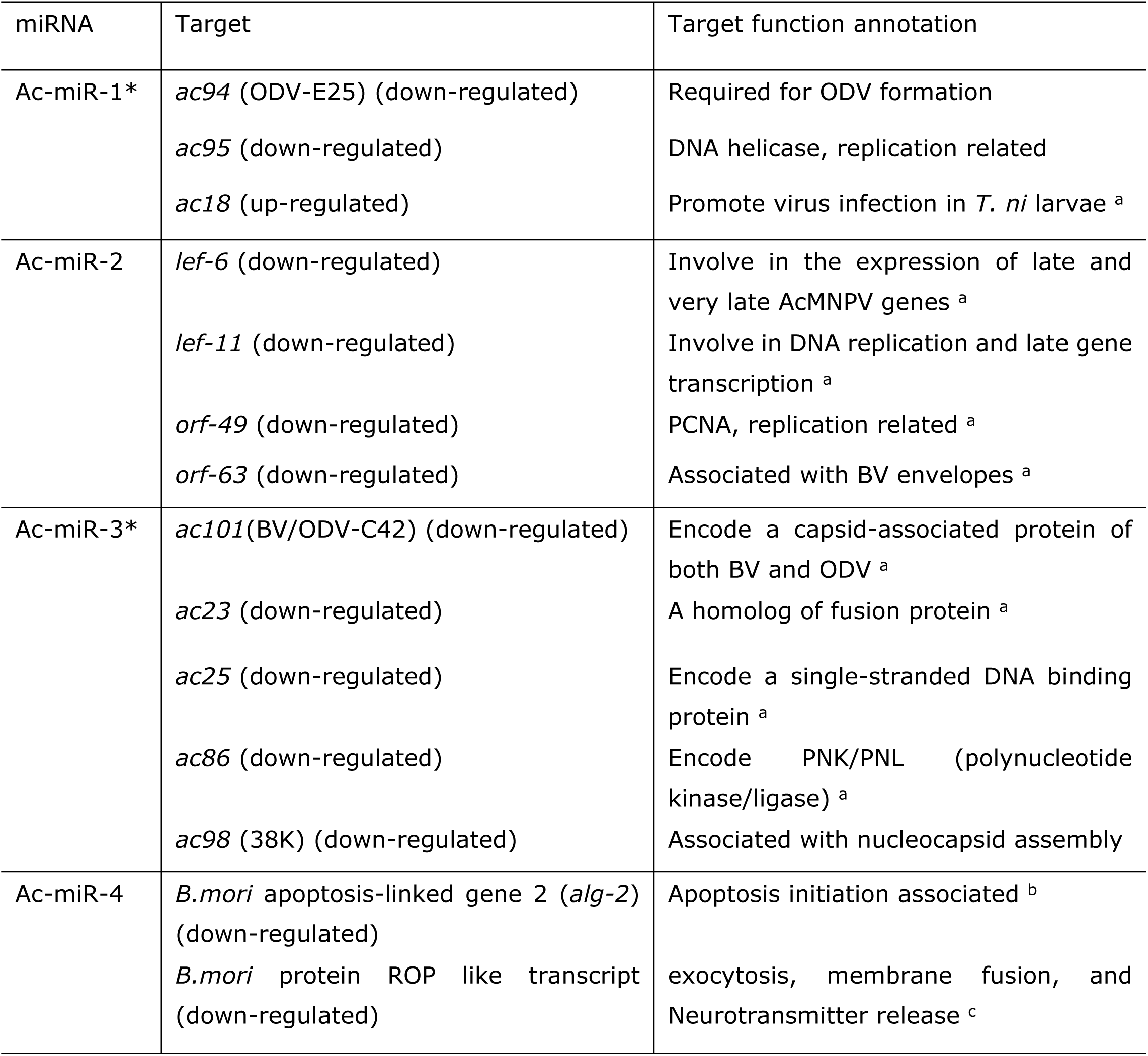
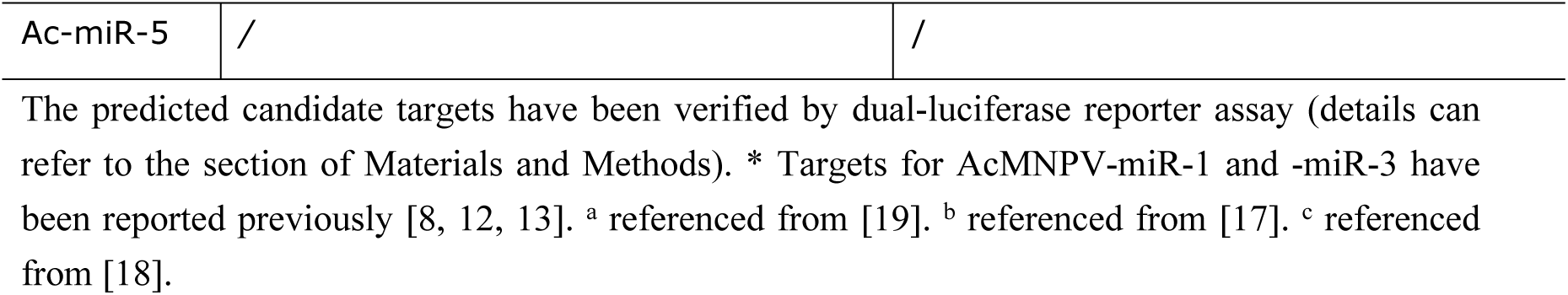
The validated primary target genes.

Five viral target genes for AcMNPV-miR-2 were predicted using the same methods described above: *lef-6, lef-11, orf-49, orf-38*, and *orf-63*. With a perfect complement to the CDS, the highest score (data not shown) was found for the late expression factor gene *lef-6* [16]. These five candidates were also tested by the dual-luciferase assay. The most distinct down-regulation was observed for *lef-6*, with *lef-11, orf-49*, and *orf-63* also showing obvious down-regulation (Table 2); in contrast, *orf-38* did not exhibit significant regulation. No potential host target genes were found for AcMNPV-miR-2.

Seven candidate targets in viral sequences were predicted for AcMNPV-miR-3. AcMNPV-miR-3 was perfectly complementary to the CDS of *ac101* (BV/ODV-C42), with an obvious inhibitory effect on this viral gene. Dual-luciferase reporter assay revealed slight down-regulation of *ac23, ac25, ac86*, and *ac98* (Table 2). No significant regulation was observed for the remainder candidates [12].

There were eight candidate targets for AcMNPV-miR-4 predicted in the viral sequence, but none of them met the criterion of a target gene after testing by the dual-luciferase reporter assay. Thirty candidate targets in the *B. mori* sequence were predicted for AcMNPV-miR-4, and all were tested by the dual-luciferase reporter assay. Among them, 14 caused up- or down-regulation to varying degrees (data not shown). We further examined *B. mori* apoptosis-linked gene 2 *(alg-2*) and the *rop* gene, and both showed down-regulation by AcMNPV-miR-4 in 293 cells (Table 2). These results were reproduced when we used the two genes of Sf9 cells. ALG-2 is involved in apoptosis initiation [17]; ROP is highly expressed in nervous system and acts as both a positive and negative modulator of neurotransmitter release. ROP is also expressed in specialized tissues in which intensive exocytic/endocytic cycles occurs [18]. Based on the two targets, we can deduce partial functions for AcMNPV-miR-4, which might include interfering with the host cell cycle, cytokine secretion, exocytosis, and membrane fusion.

Twenty predicted candidate targets each were screened using the dual-luciferase reporter assay for AcMNPV-miR-5, but none of them achieved evident regulation. Further experimental investigation for targets and function of this miRNA is still in progress.

### miRNA transcription initiation analysis

miRNA is typically generated from long primary miRNA, which is transcribed by RNA polymerase II and is rapidly cleaved to produce precursor miRNA by Drosha. Compared to the knowledge of miRNAs regulating gene expression, the understanding of miRNA transcriptional initiation is lagging far behind, mainly due to the fact that miRNA transcription start sites (TSSs) are largely unknown [20]. Addressing this issue is more complex for baculovirus miRNAs because most viral early transcripts are initiated at early promoter sequences by host cell RNA polymerase, while viral late transcripts are initiated at late promoter sequences by viral RNA polymerase. To understand how the five primary miRNAs are transcribed, we utilized sequence features derived from known baculovirus promoter motifs to scan the upstream regions of pre-miRNAs for the hallmarks of TSSs, such as early promoter consensus sequence (TATA) and late promoter consensus sequence (TAAG) [21]. To date, there is no canonical software to effectively predict the promoter sequences of miRNAs because these sequences are somewhat species specific. Nevertheless, we still employed various programs to survey the upstream sequences of the pre-miRNAs, but no effective transcription factor binding sites or obvious AT-rich regions were found.

AcMNPV-miR-1, -miR-2, and -miR-3 are all located in the antisense strand of their corresponding genes (Fig 2, Table 1). miR-1 is located in the complementary strand of *odv-e25*, and the downstream strand of *ac95* in the same orientation. TATA (80214) and three TAAG motifs were found by scanning the sequence between both the start codons of *ac95* and the precursor of miR-1. Considering miR-1 was expressed at 12 h p.i., the mapped early motif TATA unlikely initiates its transcription. TAAG (81057) was the closest one (957 nt) to miR-1 of the three late motifs, if it is an effective one, miR-1 may be transcribed from this mapped late motif. miR-2 is an early miRNA located in the complementary strand of *lef-6* and the downstream strand of the early gene *orf29* in the same orientation. TATA (24021) was found by scanning the 500 nt upstream sequence of the pre-miRNA. This early motif was located approximately 25 nt downstream of the 3’ terminus of *orf29*. Therefore, the primary AcMNPV-miR-2 may be transcribed from this closer TATA (24021) if it is not a weak motif; however, it may probably use the same TSS of *orf29* because there was only a distance of 588 nt between both start sites of *orf29* and the precursor of miR-2. Similarly, miR-3 is located in the complementary strand of *ac101* and the downstream strand of *lef-5* in the same orientation. TATA (86964) and TAAG (85941) were mapped by scanning the sequence between both the start codons of *lef-5* and the precursor of miR-3. The mapped early motif was approximately 249 nt downstream of the 3’ terminus of *lef-5*. As an early expressed miRNA, miR-3 may use this motif TATA (86964) as its transcription initiation.

The other two miRNAs, miR-4 and miR-5, are all located in the viral gene coding strand (Fig 2, Table 1). miR-4 was late expressed and located in the 5’ region of the early/late gene *cg30*. TATA (75634) and TAAG (75821) were found by scanning the 500 upstream sequence of the pre-miRNA. The two motifs were approximately -103 nt and -290 nt upstream of *cg30*; however, the TSS of *cg30* is located at approximately -242 nt [22]. Thus, miR-4 is believed to be transcribed from the TSS of *cg30*. miR-5 was located in the 5’ region of *ac-49k*, and also in the immediate early gene *ie-01* (Fig 2E), which can splice into *ie-0* [19, 22]. TATA (123610) and TAAG (123616) were found by scanning the 500 nt upstream sequence of the pre-miRNA. The two motifs are -22 nt and -16 nt upstream of the start codon of *ac-49k*. The distance between the start sites of *ie-01* and the miR-5 precursor was 943 nt. miR-5 showed distinct expression at 12 h p.i., whereas the spliced form (*ie-0*) is abundant early and comprised over 80% of the *ie-01* mRNA at 6 h p.i., but declined to 2% by 48 h p.i., and a*c-49k* transcription levels peaked at 12 h p.i [22]. Hence, miR-5 may be a splicing by-product of *ie-01* at early stage, and is more likely transcribed from the TSS of *ac-49k* at late stage of infection (Table 3).

**Table 3.** Proposed TSS of AcMNPV-encoded miRNAs.

| miRNA | Expression | Position of TATA* | Position of TAAG* | Possible TSS |
| --- | --- | --- | --- | --- |
| Ac-miR-1 | Late | 80214 | 81057 | TAAG (81057) |
| Ac-miR-2 | Early | 24021 | / | TATA (24021) |
| Ac-miR-3 | Early | 86964 | 85941 | TATA (86964) |
| Ac-miR-4 | Late | 75634 | 75821 | TSS of *cg30* |
| Ac-miR-5 | Late | 123610 | 123616 | TSS of *ac-49K* |

AcMNPV genome Accession No. NC\_001623.1. TSS: transcription start site; TATA: early motif; TAAG: late motif. \*represents they were mapped by scanning the sequence between both the start codons of situated gene or upstream gene and the precursor of miRNA or by scanning the 500 nt upstream sequence of the pre-miRNA if the size of situated gene or upstream gene is less than 500 nt.

## Discussion

Five AcMNPV encoded miRNAs were validated by experimental approaches. Three of them (AcMNPV-miR-1, -miR-2, and -miR3) are located exactly opposite the coding sequences of the corresponding viral genes and are undoubtedly the natural targets. The three miRNAs might target their perfectly complementary viral genes via RNA-mediated interference (RNAi) machinery, as do short interfering RNAs (siRNAs), cleaving the corresponding mRNAs directly [3]. The other two miRNAs are located at the 3’ (AcMNPV-miR-4) or 5’ (AcMNPV-miR-5) region of viral genes, their function pattern might be different.

AcMNPV-miR-2 and –miR-3 were detected at 3 h p.i., suggesting the two miRNAs are functional from the early stage of infection. The expression of AcMNPV-miR-1, and –miR-4 were detected at 12 h p.i., suggesting that they play roles at the late stage of infection. As for AcMNPV-miR-5, there were blotting bands showed at 3 and 6 h p.i. (Fig 3D), however, qRT-PCT detected its distinct expression at 12 h p.i. (Fig 4D). Considering this miRNA is located in both *ac-49k* and immediate early gene *ie-01*, the latter can splice into *ie-0* [19, 22]. The spliced *ie-0* is adjacent to *ac-49k* (Fig 2E). Hence, the blotted bands showed at 3 and 6 h p.i. might be splicing by-products of *ie-0/ie-01*, and miR-5 is inclined to be a late miRNA.

The majority of predicted and validated miRNA targets were found in the virus genome (Table 2), which suggests that viral miRNAs are mainly involved in active regulation of viral genes. This may include decreasing the viral DNA load and/or the number of infectious BVs to avoid host surveillance and prolong the duration of residence inside the host cell [8, 13]. In addition, most of the virus targets are late genes, which may suggest that miRNAs, especially those expressed early in infection, auto-regulate the expression levels of late genes to reduce their unnecessary accumulation early in infection [3]. AcMNPV-mir-4 is an exception, the predicted targets of which are all host genes, indicating that virus can actively invoke a sophisticated strategy to modulate host cellular physiology and biochemistry via miRNAs. These results indicate that AcMNPV-encoded miRNAs function in host-virus interactions by fine-tuning viral and/or host genes to establish infection and replicate successfully. In addition to AcMNPV-miR-4, target genes in host of other miRNAs might be identified when more complete host sequence information is available.

These AcMNPV encoded miRNAs predominantly down-regulate target genes expression. However, up-regulation also exists, such as AcMNPV-miR-1 up-regulating *ac18* [13], though it is not common. Nevertheless, down- or up-regulation of viral miRNA targets depends on the function of the target gene and serves for the ultimate aim to achieve infection. It is well known that each miRNA may have multiple targets, and one mRNA may be targeted by multiple miRNAs. The comprehensive functions of AcMNPV-encoded miRNAs rely on the complete functions of each miRNA.

Exploring TSSs of miRNAs is important for understanding how these small RNA molecules, which are known to regulate gene expression, are regulated themselves. Through sequence analysis, we deduce that miRNAs that are located in the antisense strand of viral genes may be transcribed from the mapped nearest corresponding early or late motif, while those are located in viral gene coding strands may be transcribed from the TSS of the situated gene (Table 3). This conclusion agrees with the virus economics.

The profiles of AcMNPV-encoded miRNAs appear to exhibit general characteristics of those miRNAs from baculoviruses and other insect viruses (Fig 6). Further studies on these miRNAs are on-going and will help us to gain insight into the roles of miRNAs in baculoviruses and insect-host interactions.

**Fig 6.**
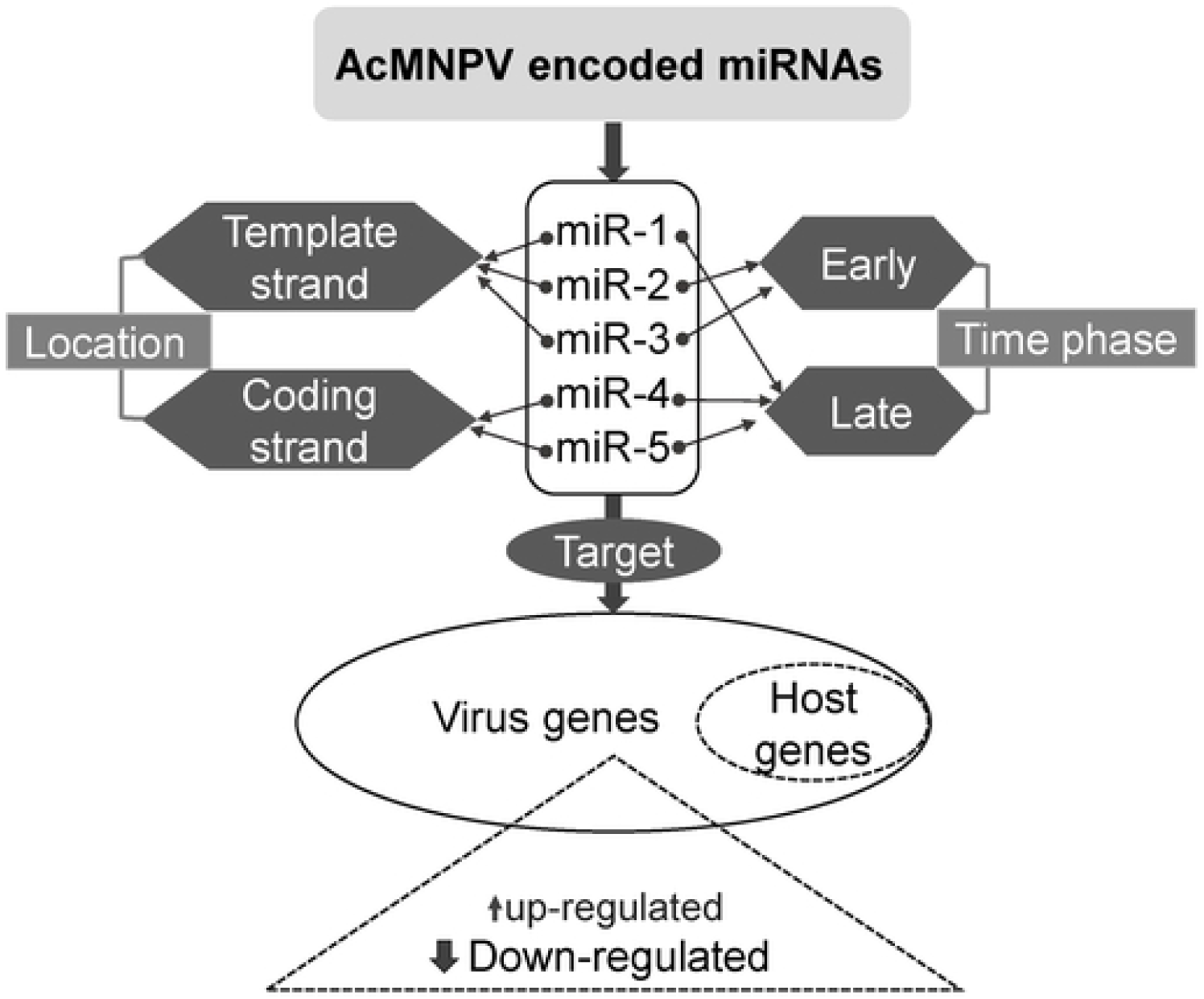
Schematic of the profiles of AcMNPV encoded miRNAs.

